# Positioning of negative feedback loops within immune signaling pathways influences gene expression noise

**DOI:** 10.1101/2024.02.22.581613

**Authors:** Danial Asgari, Ann T. Tate

## Abstract

Signaling pathways depend on negative and positive feedback loops (NFLs and PFLs) to regulate internal noise. Across diverse organisms, signaling is regulated by NFLs that function at different cellular locations. These range from NFLs functioning upstream near signal-receiving receptors to those downstream within the nucleus. While previous studies have examined the relationship between NFLs, internal noise in signaling pathways, and network topology, none have directly addressed how the cellular location of NFLs impacts noise regulation. This is significant given the almost ubiquitous presence of multi-level regulation systems within signaling pathways. Here, we use stochastic models inspired by Imd and Toll signaling to address this gap within the context of immune signaling. We use both mechanistic and evolutionary models to demonstrate how noise is regulated by NFLs and how this, in turn, affects the host’s ability to fight off infection while minimizing immunopathologic effects of excessive immune gene expression. We found that downstream NFLs reduce noise in antimicrobial peptides (AMP) expression for some parameter values. On the other hand, upstream NFLs amplify the noise, but the presence of a strong PFL can reduce this noise. Our evolutionary simulations suggest that the mechanisms through which the downstream NFL operates within the cell can affect the evolution of the upstream NFLs. The results of our study provide insight into why distinct signaling pathways are regulated by varying numbers of NFLs, which operate in different cellular locations and employ diverse mechanisms to control gene expression.

**Author Summary:** Signaling pathways are noisy biological circuits. This noise is caused by random fluctuations in the number of proteins that function within these pathways. To properly respond to external stimuli, the ratio of noise to information needs to be minimized. To regulate noise, biological pathways produce proteins that either reduce (negative feedback) or amplify (positive feedback) signaling following stimulation. Negative feedback loops can shut down signaling by interfering with the first steps of signaling, which entail the detection of stimuli. Conversely, signaling might be left intact, and instead, negative feedback loops might interfere with the last step, which is the production of output. These regulatory differences can affect noise within signaling pathways. Here, we examined this using stochastic (inherently random) models to simulate immune signaling in response to pathogens. We found that negative feedback loops that function at later stages of signaling can decrease noise in the output, while negative feedback loops that act at earlier stages amplify the noise, but this noise can be reduced by the presence of a strong positive feedback loop.

## Introduction

Organisms rely on signaling pathways to monitor environmental changes and generate appropriate responses [1–4]. Noise or stochasticity (inherent unpredictability) is a fundamental characteristic of signaling pathways [5], which originates from random fluctuations in the small number of signaling proteins and interactions between them [6]. Biological noise is influenced by many factors such as the sequence of the promoter, nucleosome occupancy, and the number of transcription factor binding sites [7]. Increased noise in gene expression is linked to cancer and aging [8,9]. Therefore, proper homeostasis demands tight control over noise.

Signaling pathways regulate noise through several mechanisms. One such mechanism involves feedback loops [10,11]. Feedback loops activate proteins that either amplify the signal (positive feedback loops, or PFLs) or attenuate it (negative feedback loops, or NFLs). The modulation of noise through feedback loops depends on several factors. These include the source of the noise, which can be either due to environmental fluctuations (extrinsic) or due to inherent fluctuations of the number of proteins involved in a signaling pathway (intrinsic) [12]. Autorepression, in which a gene suppresses its own activity, has been shown to reduce noise within signaling pathways [13,14], however, this is often linked to extrinsic, not intrinsic, noise [11,15]. The network topology and the specific mechanisms by which feedback operates can further complicate noise modulation by NFLs. For example, Chakravarty et al. showed that, in circadian networks, NFLs reduce intrinsic noise, while PFLs reduce extrinsic noise [16]. The magnitude of noise modulation also varies depending on how the feedback functions within the pathway. For example, mRNA-mediated feedback—where mRNA suppresses its own translation (e.g., microRNA)—provides better control over noise at the protein level compared to protein-mediated feedback, in which the protein downregulates the expression of its own gene [17]. In reality, many signaling pathways employ NFLs at multiple levels of the transduction cascade, which attenuate response through diverse mechanisms. However, it remains an open question whether this layering mainly provides redundancy or whether each NFL within a pathway contributes something unique or context-dependent to signal and noise control.

Immune signaling networks provide particularly salient case studies of the interaction of layered NFLs in mediating noise. The immune signaling pathways of most multicellular eukaryotes share a common architecture: signaling activates a central transcription factor that induces target genes against pathogens as well as genes encoding NFLs to turn the response back down, allowing the organism to conserve energy and mitigate immunopathological effects [18,19]. Although there are biological differences across immune signaling pathways, NFLs that regulate them generally fall into two categories: those that decrease input into the pathway by acting upstream and those that function downstream to reduce the output while maintaining the signaling process (Table 1).

**Table 1.** NFLs* at different steps of signaling in immune-related pathways. The Toll pathway in insects is only regulated by an NFL downstream of the pathway.

| Signaling pathways | Receptor | Upstream NFL | target | Downstream NFL | target |
| --- | --- | --- | --- | --- | --- |
| NF-κB (vertebrate) | TLR4 (innate)<br>TIR (innate)<br>TCR/BCR (adaptive) | A20<sup>1,2</sup> | RIP (outside the nucleus) | IκBα<sup>3</sup> | NF-κB (inside the nucleus) |
| Imd (invertebrate) | PGRP-LC | Pirk<sup>4</sup> | PGRP-LC (outside the nucleus) | Repressosome complex<sup>5</sup> | NF-κB (inside the nucleus) |
| Toll (invertebrate) | Toll | NA | NA | Cactus | NF-κB (inside the nucleus) |
| Jasmonate (plants) | JAT1 | CYP94B3<sup>6</sup> | JA-Ile (outside the nucleus) | JAM<sup>7</sup> | MYC (inside the nucleus) |

\*Here, NFLs are listed that are activated upon infection. These differ from negative regulators that control the baseline expression of genes.

Simple theoretical models that consider shared versus unique topological attributes can be used to clarify the relationship between noise, the response to stimuli, and the positioning of NFLs and PFLs within pathways while also shedding light on their evolution. For example, Asgari et al. [27] used a theoretical model of the fruit fly immune response to show that upstream and downstream NFLs contribute differently to defense against pathogens with varying proliferation rates. In the fruit fly, *Drosophila melanogaster*, the Toll pathway primarily reacts against gram-positive bacteria and fungi [28,29] while the Imd pathway responds to gram-negative bacteria [30], but both stimulate the expression of antimicrobial peptides (AMPs) through the action of NF-κB transcription factors [31]. In addition, the NF-kB activates a PFL by upregulating genes encoding receptors, which boosts the signal to provide a stronger response to the pathogen [36]. Analogously to vertebrate NF-kB signaling (Table 1), the Imd pathway includes two major NFLs: the induction of Pirk instigates the upstream removal of immune receptors from the cell surface [23] while the Repressosome acts downstream [24]. The Toll pathway, on the other hand, lacks an obvious upstream NFL and relies on the downstream Cactus protein to provide negative feedback [32,33]. In both pathways, downstream NFLs inhibit the activity of NF-κB transcription factors and reduce the output while maintaining signaling, similar to JAM transcription factors competing with MYC2 to dampen Jasmonate signaling in Arabidopsis [26]. Differences in the cellular location of NFLs (e.g., Pirk vs Repressosome) and mechanistic differences in the function of NFLs operating at the same location (Cactus vs Repressosome) could potentially yield distinct effects on noise regulation and fitness.[37].

In this study, we built stochastic models inspired by Imd and Toll signaling that contain modules representing upstream NFLs and different types of downstream NFLs. We examined how different NFLs contribute, individually and in tandem, to AMP output and noise during repeated pathogen encounters. Considering that too much immune signaling can promote pathology or impose other costs to host fitness, we conducted evolutionary simulations to examine how NFLs functioning at different locations evolve to modulate immune response, and how this, in turn, affects noise in AMP expression.

## Methods

### Models

We designed stochastic models containing only the fundamental components of an immune signaling pathway. These entail a receptor that detects the pathogen and activates a central transcription factor, which in turn induces AMPs that kill the pathogen, along with NFLs that attenuate the response. Inspired by the Imd pathway, we included two NFL modules, acting upstream (Pirk that removes receptors) and downstream (Repressosome that competes with the central transcription factor Relish over binding to the promoter of AMP genes). Inspired by Toll topology, we supplied a cactus-like module that could be swapped for the Imd repressosome module to provide an alternative downstream NFL. Unlike the repressosome, cactus inhibits the response by degrading Relish before it enters the nucleus (Fig.1). We implemented the two models using the Gillespie algorithm [38], where reactions capture the change of proteins within an immune signaling pathway. At each step, a reaction is probabilistically chosen and carried out. The likelihood of a reaction depends on its rate relative to the total rates of all reactions. In both models, time is discrete (T = 1,000). The complete models are presented in detail in the S1 text. Both models entail two sets of reaction rates: reaction rates for bacteria and reaction rates controlling immune signaling. If both sets are empty (i.e., all reaction rates are 0), no reactions occur. If both sets contain non-zero rates, two reactions occur in one step (one for bacteria, which is either elimination or proliferation, and one for a randomly chosen protein of the immune network). Otherwise, one reaction occurs.

**Fig. 1.**
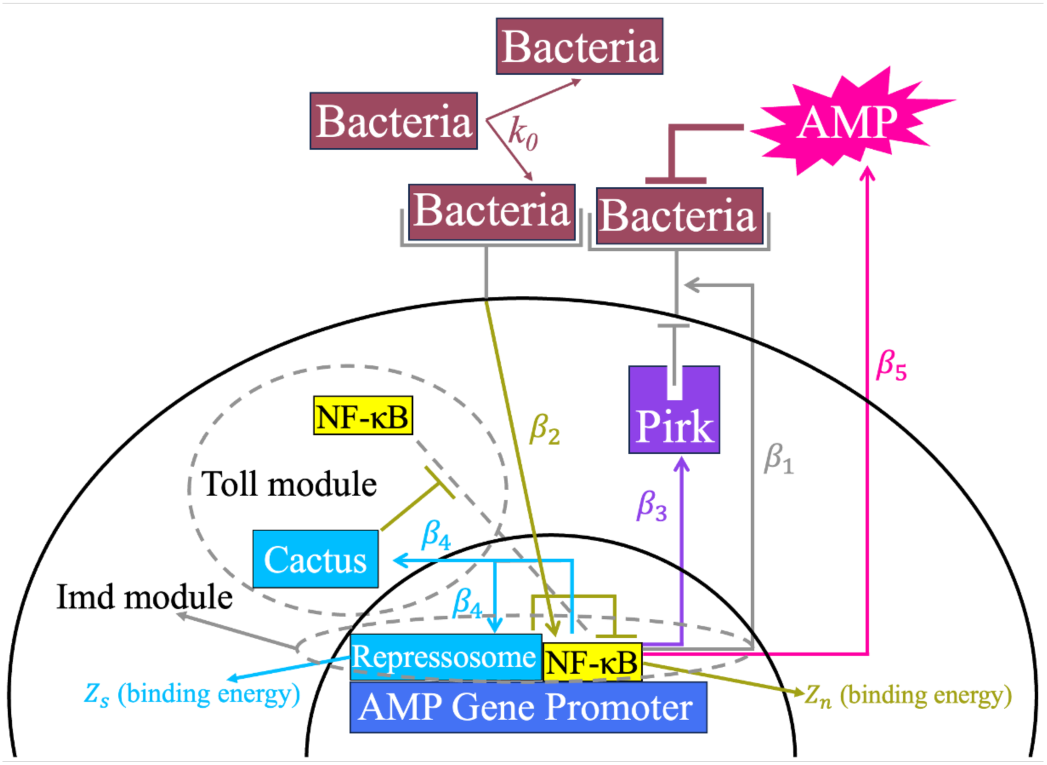
Simplified diagrams of models inspired by the Imd and Toll pathway. Bacteria proliferate at a rate *k_0_* and bind to receptors. NF-κB transcription factors are activated by bacteria-receptor complexes at a rate β_2_. NF-κB induces the production of receptors (β_1_), Pirk, which is the upstream NFL(β_3_), Repressosome (in Imd-like model), and Cactus (in Toll-like model), which are the downstream NFLs (β_4_), and AMPs (β_5_). Pirk reduces receptors in both models. Repressosome competes with NF-κB for binding to the promoter of the AMP gene, and Cactus inhibits the entry of NF-κB to the nucleus. The binding energies of the NF-κB and Repressosome are *Z*_*n*_ and *Z*_*s*_, respectively. The colors of the arrows are based on the affected components in each reaction.

The stochastic reactions described above simulate inherent noise within the Imd and Toll-inspired systems. To understand how these systems function in the absence of noise, we developed deterministic models of the systems described above (S1 Text) using systems of ordinary differential equations (ODEs). To solve these equations, we employed the Backward Differentiation Formula (BDF) method via the solve_ivp function in Python, setting atol=10^−6^ and rtol=10^−8^ to ensure numerical accuracy.

### Parameter regimes

We considered four parameter regimes to examine the effect of NFLs on the dynamics of Imd and Toll-inspired models (Table 2). The four regimes are based on two parameters that specify the production rate of NFLs. The upstream (**β**_3_) and downstream (**β**_4_) NFLs are either highly expressed (β_3_ or β_4_ = **10**) or knocked down (β_3_ or β_4_ = **0**). We chose these values based on the qualitative assessments of several simulations before conducting the systematic analysis of the models. We conduct additional simulations for other values of β_3_ and β_4_ for specific scenarios as discussed in the Results section.

**Table 2.** Four parameter regimes capturing different NFL activities upstream and downstream of Imd and Toll signaling

| Regime | parameters |
| --- | --- |
| Both knocked down | $\beta_3$ and $\beta_4 = 0$ |
| Upstream active | $\beta_3 = 10$ ; $\beta_4 = 0$ |
| Downstream active | $\beta_3 = 0$ ; $\beta_4 = 10$ |
| Both active | $\beta_3$ and $\beta_4 = 10$ |

There are four background parameters (β_1_, β_2_, β_5_, λ) in Imd and Toll-like models. These are distinct from parameters that determine the dynamics of NFLs (β_3_, β_4_, *Z*_*s*_) and the binding energy of NF-κB (*Z*_*n*_). For each of the background parameters, we considered 10 different values. The background parameter values range from 1 to 10 for β_1_, β_2_, β_5_ and from 0.1 to 1 for λ. We chose smaller values for λ because larger values result in a highly inefficient immune response (i.e., the immune proteins degrade before the transmission of the signal). This constitutes a total of 10,000 simulations to explore all combinations of background parameters for one parameter regime (10,000 x 4 = 40,000 parameters for the four regimes in Table 2). In the Imd-like model, across all parameter regimes, we maintained the following condition: *Z*_*s*_ = *Z*_*n*_ = 1, unless otherwise stated. This ensures that the binding of the Repressosome is only influenced by the relative number of NF-κB to Repressosome, which in turn is influenced by β_4_ that varies across the four regimes. For consistency in the Toll-like model, we set *Z*_*n*_ = 1.

### Calculation of AMP noise, average AMP expression, and fitness

For each parameter set, we conducted 10,000 replicate simulations, and quantified noise in AMP expression at every time point of the host life time (T = 1,000) by calculating the coefficient of variation [39] (Eq. 1). We then determined the magnitude of the noise by calculating the arithmetic average of the noise across all time points (Eq. 2). We measured the average AMP expression for each parameter set by calculating the arithmetic mean of average AMP expression (average across 10,000 replicate simulations) across the lifetime of the host (T = 1,000) (Eq. 3).

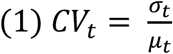

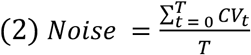

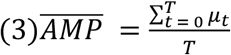

Here μ*_t_* and σ_*t*_ are the average and standard deviation across 10,000 replicate simulations. If μ*_t_* = σ*_t_* = 0, *CV_t_* is set to 0.

We estimate the fitness by either calculating the host’s ability to reduce bacterial load (Eq.4) or by the host’s ability to fight off infection while minimizing the AMP expression (Eq.5). Eq. 4 reflects a scenario where the pathogen virulence far outweighs immunopathology, while Eq. 5 weights them more equally. In Eqs. 4 and 5, *A* and *B* are the average value for these variables across the lifespan of the host. Eq.5 is used to run evolutionary simulations of Imd and Toll signaling.

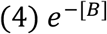

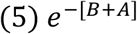

Repeated encounters with pathogens create strong evolutionary pressure, which drives the evolution of an optimized immune signaling network. Here, for all simulations, the host encounters bacteria periodically at every 200 time points within the host lifetime (T = 1000, five total encounters during the host lifetime). This captures the real-life condition of multiple encounters with the pathogen [40,41].

### Evolutionary simulations

We ran evolutionary simulations to find optimum parameters controlling the expression rates of NFLs upstream and downstream of both Imd and Toll-like models. Each evolutionary simulation consists of 500 replicates, each starting with a random set of parameter values, and lasting for 5,000 generations. We are interested in the final distribution of parameters at the end of each simulation. At each generation, a single mutation happens in a randomly selected parameter among six parameters, which include four background parameters and two that regulate upstream and downstream NFLs. The mutation changes the parameter value by either increasing or decreasing it by 1 unit with equal probability, except for the degradation rate, which changes by 0.1 unit. The boundary condition keeps parameters within the range [0.1,1] for the degradation rate and between [1,10] for other parameters. If the mutation increases fitness estimated using Eq.5, the mutation is accepted, otherwise, it’s rejected. The simulations do not consist of populations of evolving individuals; rather, they entail changes in parameter values to optimize the immune response to pathogens according to Eq.5.

## Results

### The downstream NFL can suppress noise in gene expression, while the upstream NFL reduces noise through its interaction with the PFL

Here, we use models inspired by Imd and Toll signaling pathways to understand the effect of NFLs on noise in the final product of immune signaling (i.e., AMP) and ultimately host fitness. Our model has parameters controlling the strength of NFLs, and background parameters that specify other reactions, such as NF-κB activation rate, AMP expression rate, and degradation rates of signaling proteins. The upstream NFL in both models is Pirk, which removes cell surface receptors. The downstream NFL in the Imd-like model is the Repressosome complex, which antagonistically competes with NF-κB for binding to the promoter of the AMP gene. The downstream NFL in Toll-like model is Cactus, which prevents entry of NF-κB to the nucleus.

To understand how upstream and downstream NFLs change AMP noise, for both models, we compared noise in AMP expression in a circuit without NFLs (i.e., both knocked down) to one with an upstream NFL, a downstream NFL, or both (Table.1). This yields three pair-wise noise level comparisons (Δ noise). To avoid confounding noise (coefficient of variation) with the average AMP expression, we only compared noise levels between background parameter sets with the same average AMP expression, following Borri et al. (2016) [42]. We found that in both models the upstream NFL amplifies the noise (Fig.2A). On the other hand, the downstream NFL suppresses noise in AMP expression, but only for some background parameter values (Fig.2B). When both NFLs are active, the effect of the upstream NFL (noise amplification) is stronger than the downstream NFL (Fig.2C). For the Toll-like model and the stronger binding of the downstream NFL in the Imd-like model, the downstream NFLs suppress noise only when AMP expression is small (second row in Fig.2B and Fig.S1).

**Fig. 2.**
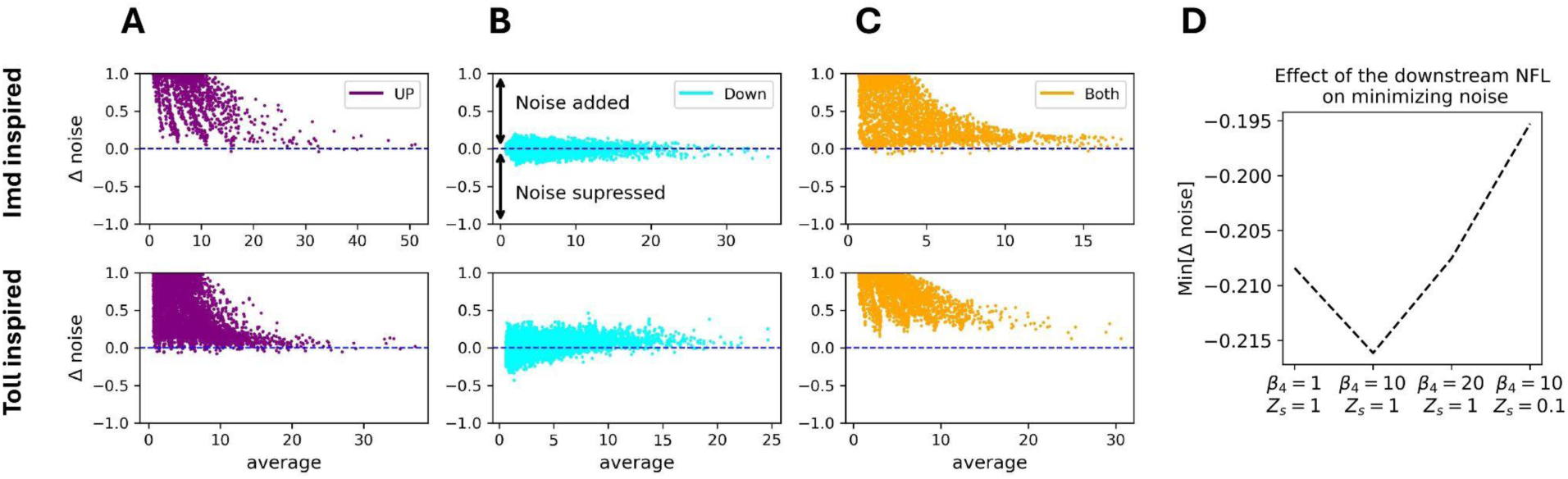
The downstream NFL suppresses noise (Δ noise < 0) in AMP expression for some background parameters, while the upstream NFL amplifies noise for both Imd and Toll-like models. Here, the Y-axis shows the difference in AMP noise between a signaling pathway without NFLs (i.e., both NFLs are knocked down) and one with one NFL (first two columns to the left) or both (third column to the right) NFLs, when the average AMP expression between the two circuits (X-axis) are almost identical (|Δ average| ≤ 0.001). The first row corresponds to the Imd-inspired model, while the second row corresponds to the Toll-inspired model. The Y-axis is tanh normalized. Here, the bacterial proliferation rate is low (*k*_0_ = 0.1). For *k*_0_ = 0.5, see Fig.S2.

Lestas et al. (2010) [43] used information theory to show that noise suppression by negative feedback has a fundamental limit. Therefore, we explored the limit of noise suppression by calculating min[Δ noise] for different strengths of the downstream NFL. To this end, we measured min[Δ noise] for weak (β_4_ = 1), strong (β_4_= 10), and very strong (β_4_=20) expression of the downstream NFL, as well as for strong expression with 10× stronger binding to the promoter (β_4_=10 and *Z*_*s*_ = 0.1). In all other cases, binding strength was kept at *Z*_*s*_ = 1. We found that maximum noise suppression occurs at strong downstream NFL expression (β_4_= 10), while both lower and higher levels result in a weaker suppression (Fig.2D).

In biological circuits, NFLs and PFLs interact to adjust the level of noise [10]. Therefore, we explored how the presence of one or both NFLs influences AMP noise across varying levels of PFL. In both models, the receptor that detects the pathogen is activated by its downstream protein (NF-κB) as a PFL. For the four parameter regimes, we plotted AMP noise as a function of receptor activation rate (receptor activation ranging from 1 to 10) (Fig.3A), and calculated the slope of the resulting line, while keeping other background parameters (other than receptor activation rate) fixed. We did this for 1,000 different combinations of fixed background parameter values. Unlike previous analyses, here we focus on changes in noise within each regime for different receptor activation rates, whereas before (Fig.2) we compared the noise of each regime to a regime with no NFLs. We observed that, for most parameter values, noise is reduced for higher levels of PFL when both NFLs are present (Fig.3B, bottom right panel).

**Fig. 3.**
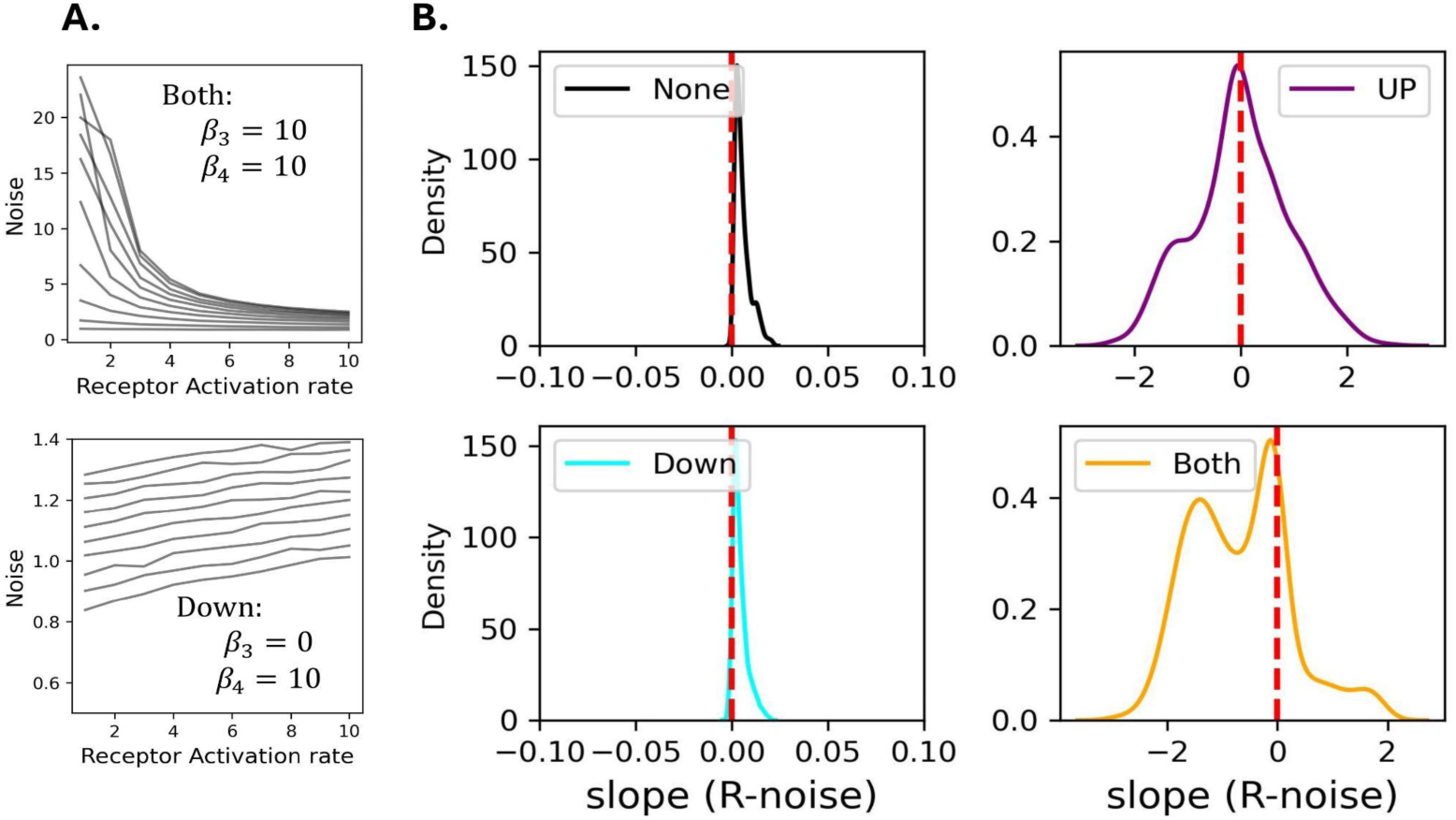
Within each regime, the noise in AMP expression is reduced for higher intensity of the PFL when both NFLs are highly expressed, primarily due to the upstream NFL. Panel A shows 10 trajectories of noise (y-axis) against receptor activation rates (x-axis) within two regimes: both NFLs active (top) and only the downstream NFL active (bottom). In Panel B, the X-axis shows the slope of the line between the strength of the PFL (R activation rate), which ranges from 1 to 10 (x-axis in Panel A), and noise in AMP expression (y-axis in Panel A). When the slope is negative, AMP noise is reduced for higher rates of receptor activation. The results are shown for Imd signaling and *k*_0_ = 0.1.

When only the upstream NFL is present, noise is reduced for some parameter values and increased for others, while it is almost unchanged when only the downstream NFL is present (Fig.3B). For the Toll-inspired model we observed the same results except that when only the upstream NFL is present noise is almost always reduced (Fig.S3). Overall, when both NFLs are active, the presence of a strong PFL reduces noise, but this effect is mostly linked to the presence of the upstream NFL.

We next explored the effect of increasing NF-κB activation rate (β_2_) rather than the receptor which activates NF-κB as a PFL (i.e., focusing on positive regulator rather than the positive feedback loop) on AMP noise within each of the four regimes. The results closely mirrored those observed when changing the PFL (Fig.S4). It is worth noting that here we do not control for the average expression when comparing noise levels. Thus, the reduction of noise could be partly due to an increase in gene expression. However, the key observation is that amplifying the signal, whether through the PFL (i.e., the receptor) or the positive regulator of the signaling pathway (NF-κB) reduces noise, especially when both NFLs are present, and this effect is mostly linked to the presence of the upstream NFL.

### Both NFLs evolve for the Imd-inspired model, while the Toll-inspired model only evolves toward a downstream NFL

To understand the effect of AMP noise on fitness, we first evaluated the effect of AMP noise on the organism’s ability to fight off infection. To this end, we estimated fitness based on the bacterial load (*F* = *e*^−*B*^; as *B* → ∞, *F* → 0). We compared host fitness under conditions with (via Gillespie simulation) and without noise (via ODE simulation). For optimal parameters (i.e., high fitness for the ODE system—that is, in the absence of noise), we observed that the fitness from the Gillespie algorithm highly depends on the AMP noise level. Specifically, optimal parameters with high noise have low fitness, while those with low noise have high fitness (Fig.S5). This shows that noise in AMP expression impairs the host’s ability to fight off infection.

Given that noise is regulated differently by upstream and downstream NFLs (Fig.2 and Fig.3) and that noisy AMP expression impairs the host response to the pathogen (Fig.S5), we next asked how these NFLs evolve to optimize host fitness. To this end, we conducted evolutionary simulations starting with random parameters that mutate by adding or subtracting a small value to one of the six evolving parameters. These include background parameters and parameters controlling the expression levels of the two NFLs. We assumed that the fitness depends on the host’s ability to reduce bacterial load while minimizing AMP expression to avoid immunopathology (*e*^−[*A+B*]^). Under this scenario, NFLs are expected to evolve and optimize the level of AMP expression. We observed that when the downstream NFL competes with NF-κB (Imd-inspired model), most simulations evolved toward either a strong downstream NFL (and a weak upstream NFL) or toward a circuit with a strong expression of both NFLs. On the other hand, when the downstream NFL affects the translocation of NF-κB (Toll-inspired model), the system evolved only toward a strong downstream NFL (Fig.4A). This is interesting given that Toll signaling in *D. melanogaster* lacks an upstream NFL. For both models, in some simulations, AMP noise increased during evolution (red stars in Fig.4A). These results were robust to changes in the bacterial proliferation rate (Fig.S6). When we used *e*^−[*B*]^ as the fitness function, noise did not evolve. This suggests that the evolution of noise is driven by the signaling pathway’s attempt to lower average AMP expression, which minimizes energetic costs and immunopathology (Fig.S7).

**Fig. 4.**
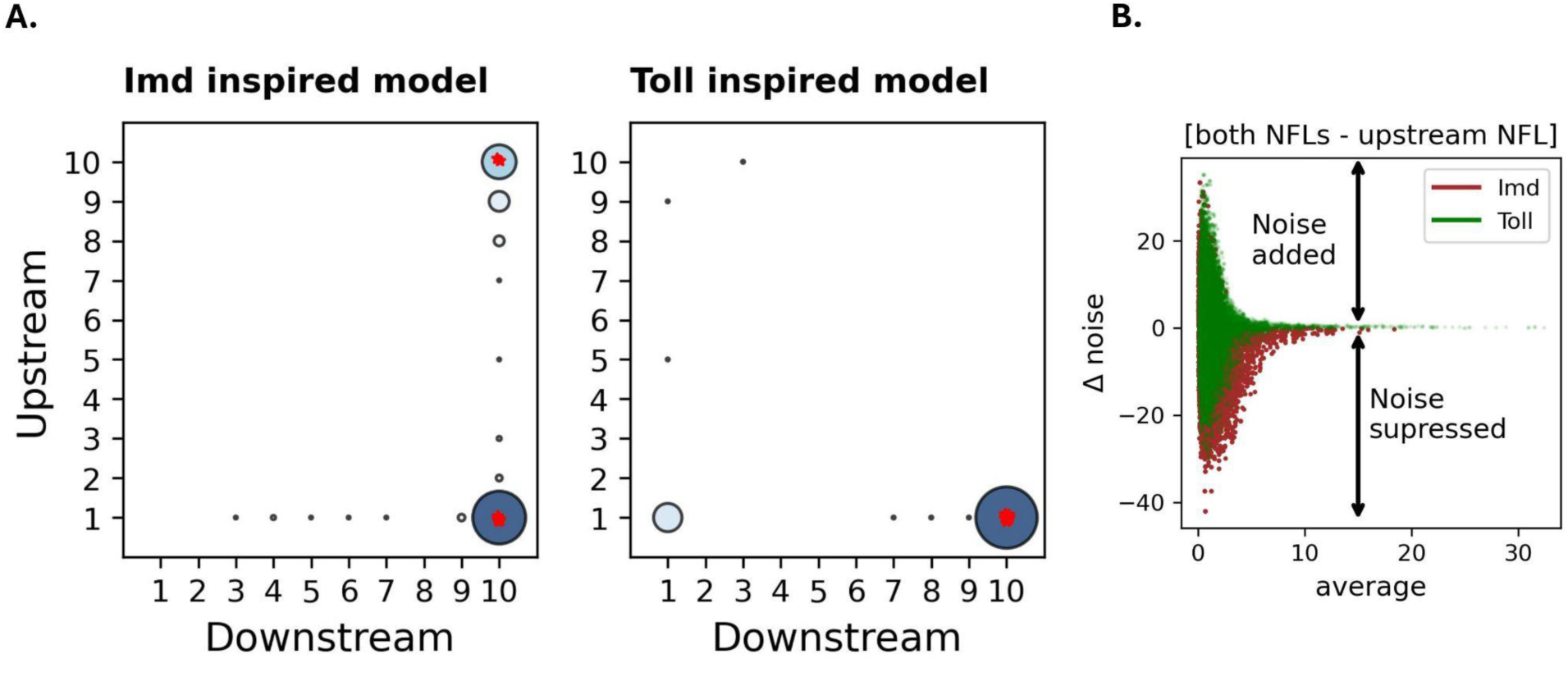
Imd signaling evolves toward a system with a downstream NFL or a system regulated by both NFLs, while Toll solely evolves toward a system with a downstream NFL. Panel A shows the final values of parameters controlling downstream (X-axis) and upstream (Y-axis) NFLs after 5,000 generations. Stars show simulations in which noise increases across generations. The size of the circles and the darkness of the color for every combination of NFLs are determined by the proportion of simulations that converge to that combination of NFLs. Panel B shows the difference in noise values between average-matched parameters across two regimes: both NFLs vs just upstream for Toll and Imd signaling.

Because the only difference between Toll and Imd-inspired models is in the mechanism of action of the downstream NFL, we hypothesized that compared to Toll-inspired model, the downstream NFL in Imd-inspired model is more effective in reducing the noise, which is amplified by the upstream NFL (Fig.2). To test this, we asked how the presence of both NFL in the two models affect the noise compared to a circuit with only the upstream NFL. To this end, we compared AMP noise between average-matched parameters across these two regimes (Fig.4B). We observed that the Imd-inspired model is slightly more effective in suppressing noise than Toll signaling. This could partly explain the evolution of both NFLs in the Imd-inspired model.

To further explore the effect of AMP noise on the evolution of NFLs, we ran deterministic simulations of both Imd and Toll signaling for two bacterial proliferation rates (*k*_0_ = 0.1,0.5), while allowing the six parameters to evolve. We observed that both signaling networks evolve toward a broader range of parameter values (Fig.S8). This is because for most parameters, ODE has a high fitness due to its rapid response to the pathogen. However, for a lower bacterial proliferation rate, both signaling pathways evolved toward an upstream-biased NFL, consistent with the results from Asgari et al. [27]. The evolution of the upstream-biased signaling pathways in deterministic but not stochastic simulations confirms the deleterious effect of the upstream NFL caused by high noise in AMP expression (Fig.2), which should be more pronounced for the Toll-like model (Fig.4B).

## Discussion

We estimated the impact of upstream and downstream NFLs on the noise level in AMP expression using stochastic models inspired by Imd and Toll signaling. We found that, compared to a circuit with no NFL, the upstream NFL amplifies noise, whereas, for some parameter values, the downstream NFL reduces noise (Fig.2). Although the upstream NFL amplifies noise, this noise can be reduced by the presence of a strong PFL, likely due to a concomitant increase in gene expression (Fig.3). This is important given that noisy AMP expression reduces the host’s ability to fight off infection (Fig.S5). We have also shown that the noisy expression of AMP evolves when AMP expression incurs a cost and needs to be minimized (Fig.4A and Fig.S7). The evolutionary pattern of NFLs was consistent with the structures of Imd and Toll signaling in *D. melanogaster* (Fig.4A). Specifically, we found that the Imd-inspired model evolves toward both upstream and downstream NFLs, whereas the Toll-inspired model only converges toward the downstream NFLs. We suggest that this is due to the effectiveness of the downstream NFL within the Imd-inspired model in reducing noise (Fig.4B).

The observation that an upstream NFL amplifies noise makes intuitive sense because the upstream NFL reduces the input to the signaling pathway (e.g., by reducing receptors that receive environmental signals). This stops the signaling before it propagates, which significantly reduces the expression of the target gene (here AMP). A low expression results in significant fluctuations in gene expression. On the other hand, a downstream NFL keeps signaling intact but only regulates the target gene expression either directly (Repressosome in Imd) or indirectly (Cactus in Toll). Borri et al. (2016) [42] showed that stronger protein-promoter binding in a negative feedback system reduces noise in enzymatic reactions. This is consistent with the effect of the downstream NFL on noise reduction in Imd signaling, where the Repressosome binds to the promoter of AMP as an NFL. Because we observed a similar effect for the downstream NFL in the Toll-inspired model, where the downstream NFL does not bind to the promoter but instead reduces transcription factor entry into the nucleus, we suggest that downstream NFLs, regardless of their mechanism of action, are capable of reducing noise in biological circuits for at least some parameter values.

Signaling networks usually take advantage of both positive and negative feedback loops to provide an appropriate response to environmental changes [10]. For example, a PFL can contribute to plasticity by inducing a switch-like behavior [44] or promoting oscillations [45]. Here we identify another interesting role for PFLs in biological circuits. We show that a strong PFL can reduce the noise induced by the presence of an upstream NFL. In the Imd signaling pathway, Pirk (or PGRP-LB) reduces the input to the system by preventing an interaction between the bacterial peptidoglycan and the receptor [23,46]. This, according to the predictions of our model, could increase noise in AMP expression, which reduces host’s ability to fight off infection, even compared to the same level of AMP expression without noise. NF-κB, however, not only activates Pirk and PGRP-LB but also upregulates the receptor production itself [37]. This serves as a PFL, which attenuates noise in AMP expression and possibly improves the host’s ability to respond to pathogens.

Signaling pathways show variation in both the number of NFLs and in the mechanism by which NFLs regulate signaling. For example, Wnt signaling, which is conserved across multicellular organisms and controls different aspects of cellular development [47], is controlled by both upstream (DKK3) and downstream (AXIN2) NFLs [48,49]. Hippo signaling, which is also crucial for the developmental process across species, is regulated by downstream NFLs (LATS1/2), which interact with transcription factors that activate target genes [50,51]. The interaction between an NFL and its target protein can be direct or indirect. For example, in NF-κB signaling of mammals, A20 indirectly interacts with the receptor (through RIP-1), whereas Pirk in Imd signaling directly interacts with the receptor. These differences in the number and function of NFLs can have consequences on the evolution of NFLs. For example, Asgari and Tate [52] have shown that downstream NFLs are more resistant to evolutionary changes due to direct interaction with the target genes. Here, by incorporating noise in our model we show that t the downstream NFL in Imd signaling (Repressosome), which directly binds to the promoter of the target gene, is more efficient in noise reduction than the downstream NFL in Toll signaling (Cactus), which only inhibits the entry of the transcription factor to the nucleus. Given that we found upstream NFLs amplify noise (Fig.2), this could partly explain why Toll signaling in organisms such as *D. melanogaster* lacks an upstream NFL.

In conclusion, by using stochastic models inspired by the two main immune signaling networks in invertebrates, we estimate the effect of upstream and downstream NFLs on noise and ultimately on host fitness. Our research provides insights into why different signaling pathways vary in the number and placement of upstream and downstream NFLs (and PFLs), helping us better understand and appreciate the diversity of signaling architectures.

## Supporting information

Supplemental file for the manuscript titled Positioning of negative feedback loops within immune signaling pathways influences gene expression noise

## Acknowledgments

This work was supported by the National Institute of General Medical Sciences at the National Institutes of Health (grant number R35GM138007 to A.T.T.).

This work used Purdue Anvil resources at Purdue University through allocation BIO240182 from the Advanced Cyberinfrastructure Coordination Ecosystem: Services & Support (ACCESS) program, which is supported by U.S. National Science Foundation grants #2138259, #2138286, #2138307, #2137603, and #2138296.

## Author Contributions

D.A. and A.T.T. conceived and designed the study. D.A. performed the simulations and wrote the first draft. D.A. and A.T.T. contributed to the final draft.

## Data Availability Statement

We performed all simulations in Python. All codes are accessible at: https://github.com/danialasg74/Positioning-of-negative-feedback-loops-and-noise/tree/main

## Supporting information

**S1 Text. Supplementary information file, including a detailed description of the Gillespie algorithm and the deterministic model.** The first section contains a detailed description of simulations of Imd and Toll signaling using the Gillespie algorithm. The second section shows the system of ODE modeling deterministic signaling pathways.

**Fig.S1.**
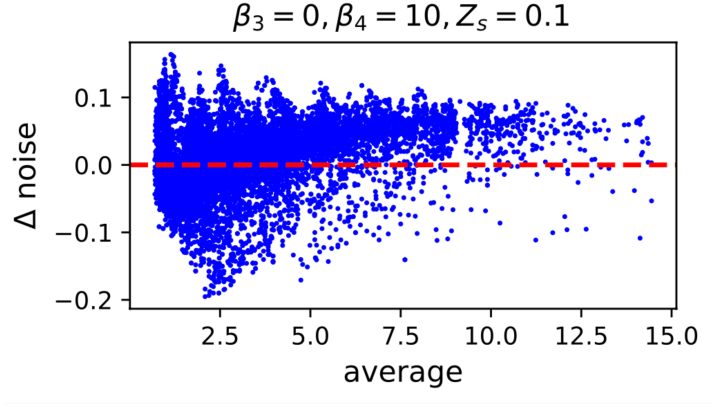
In Imd signaling, upon strong binding of the downstream NFL to the promoter of AMP genes, AMP noise is suppressed only when gene expression is small. The change in noise is shown for a downstream biased regime with a strong expression of the downstream NFL (β_4_ = 10) and 10x stronger binding (*Z*_*s*_ = 0.1) compared to Fig.2 is compared to a circuit with no NFL. The comparison is done for parameters with the same average AMP expression.

**Fig.S2.**
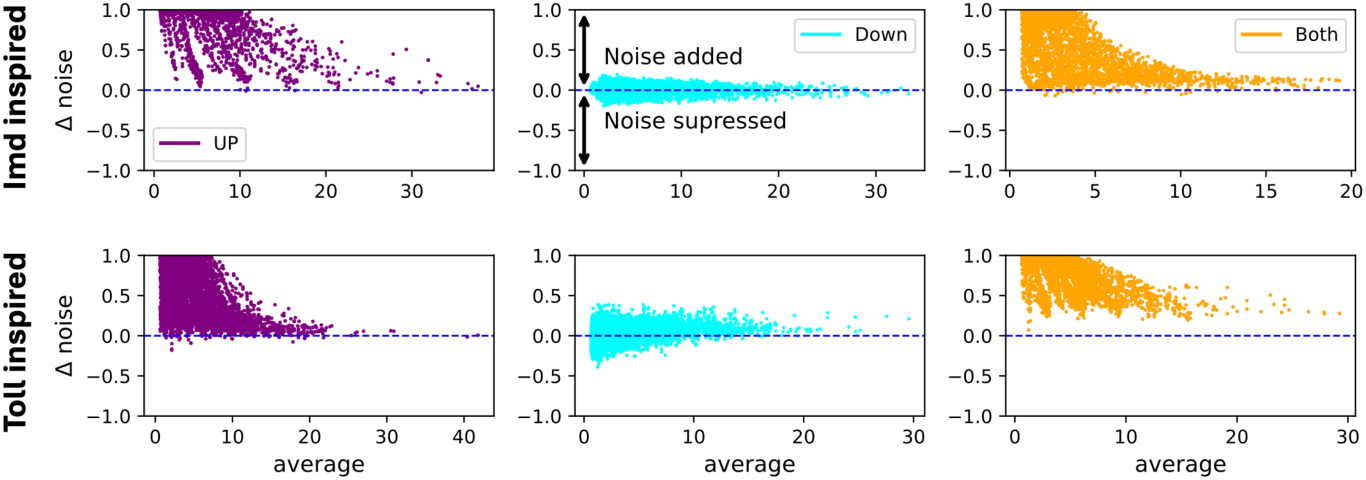
For a high bacterial proliferation rate, the downstream NFL reduces AMP noise for some parameters, while the upstream NFL amplifies noise. The change in noise across three regimes (upstream biased, downstream biased, both from left to right) compared to a circuit with no NFLs. The comparison is done for parameters with the same average AMP expression. Here, the bacterial proliferation is set to *k*_0_ =0.5.

**Fig.S3.**
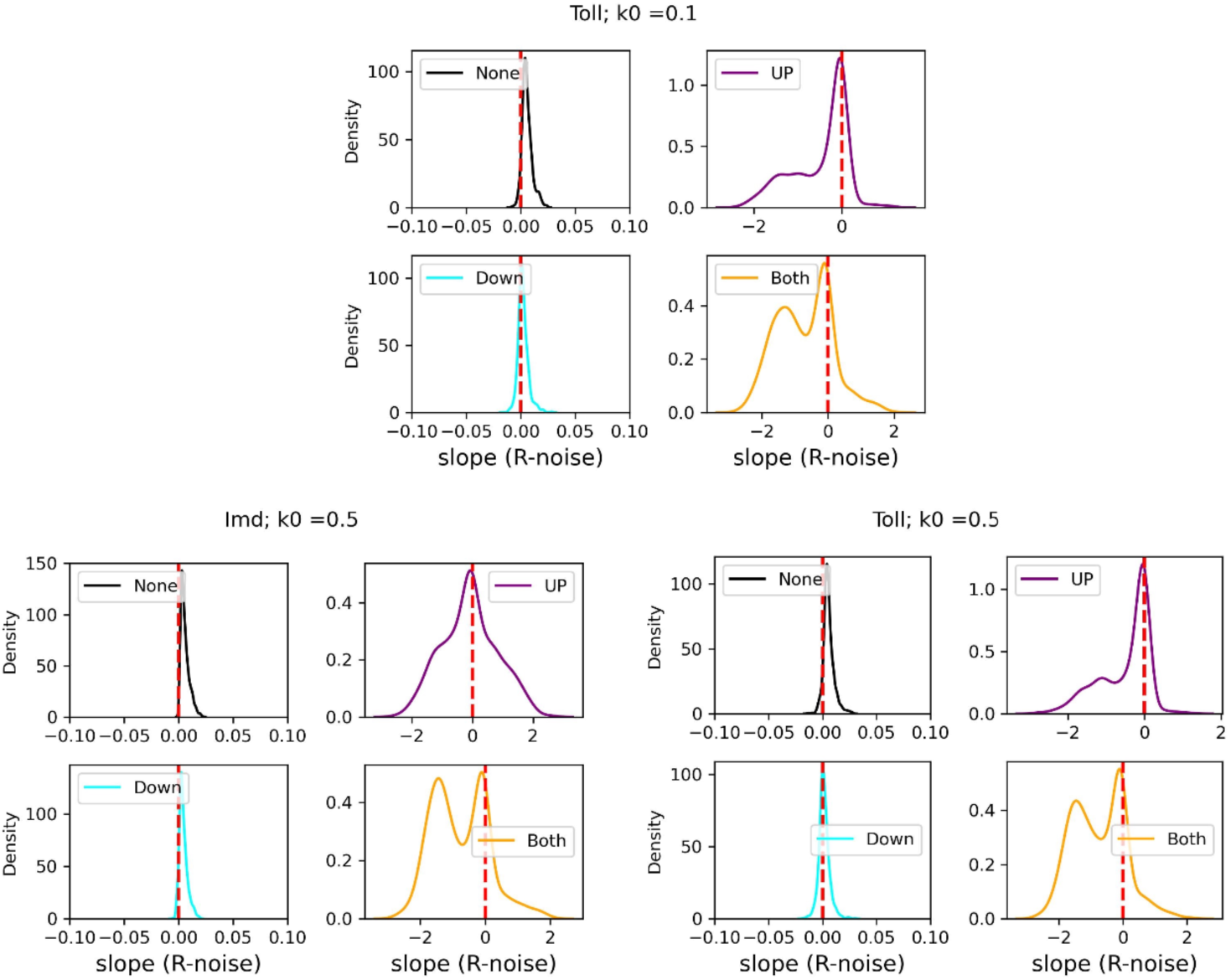
The combination of the upstream NFL and a strong expression of PFL reduces AMP noise. The slope between receptor activation rate and AMP noise is plotted for Toll signaling (*k*_0_ = 0.1) at the top, Imd signaling (*k*_0_ = 0.5) at the bottom left, and Toll (*k*_0_ = 0.5) at the bottom right. Positive slope indicates that increasing receptor activation rate increases noise, whereas negative values indicate that increasing the receptor activation rate reduces noise.

**Fig.S4.**
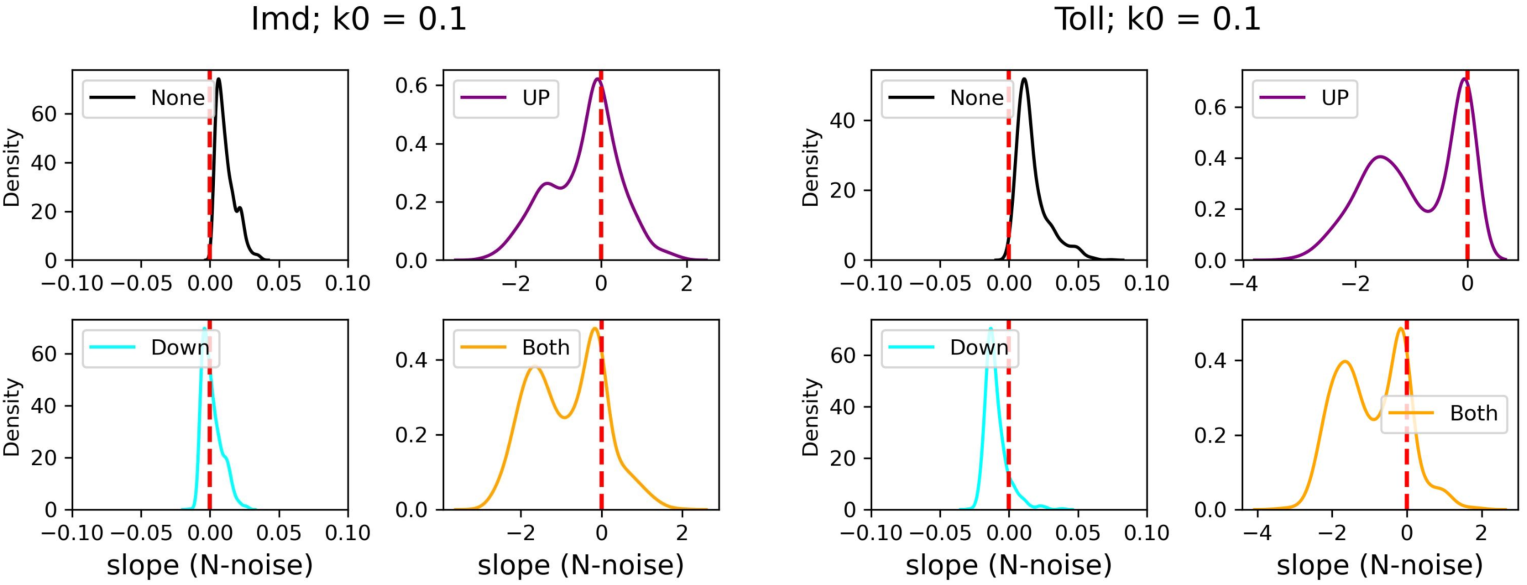
The combination of the upstream NFL and a strong expression of the positive regulator (NF-κB) reduces AMP noise. The slope between NF-kB activation rate and noise is plotted for Imd signaling on the left and Toll signaling on the right. Positive slope indicates that increasing NFkB activation rate increases noise, whereas negative values indicate that increasing NFkB activation rate reduces noise. Here, we set *k*_0_ = 0.1.

**Fig.S5.**
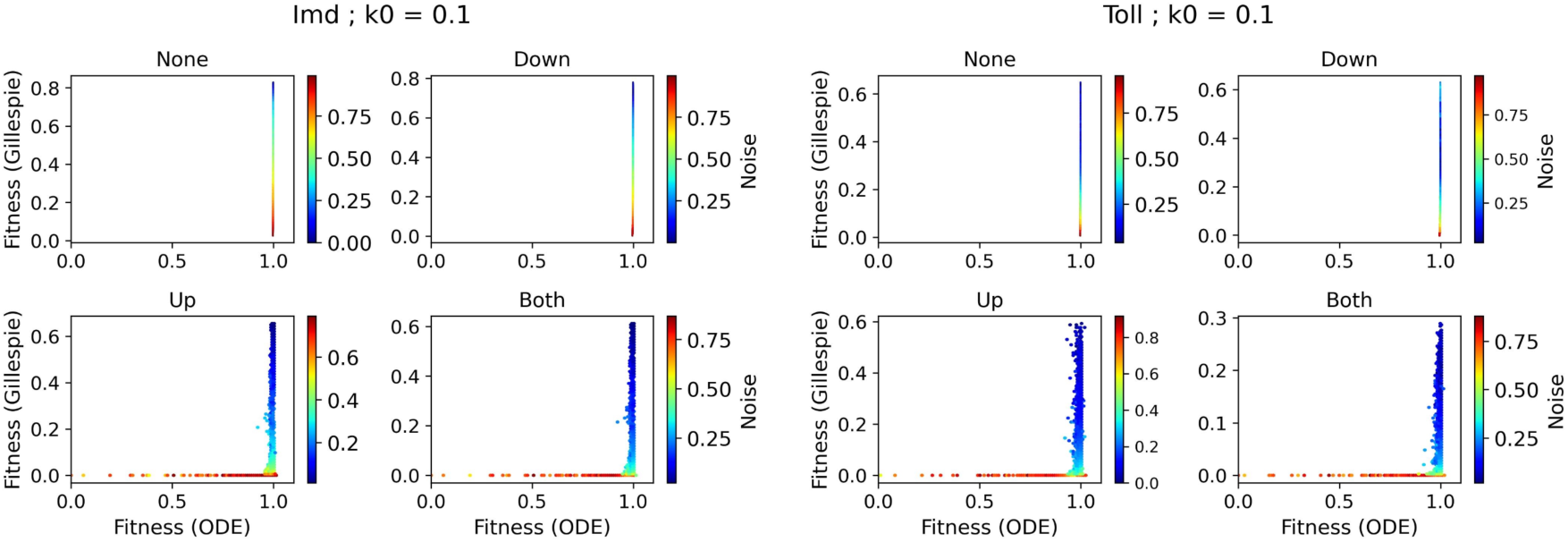
Noise reduces fitness even for optimal parameters (high fitness in the ODE model). Fitness (*e*^−*B*^) values for different combinations of background parameters are shown using the Gillespie algorithm (Y-axis) and the ODE system (X-axis). The heatmap color represents the level of noise. Four parameter regimes are shown: No NFL, downstream NFL only (Down), upstream NFL only (Up), and both NFLs. To enhance the visibility of the pattern, the noise values are log-normalized. Imd signaling results are shown on the left, and the results for Toll signaling are on the right. For each pathway, in the top two panels, the fitness of the ODE system is always near 1, regardless of the background parameter value. Here we set *k*_0_ = 0.1.

**Fig.S6.**
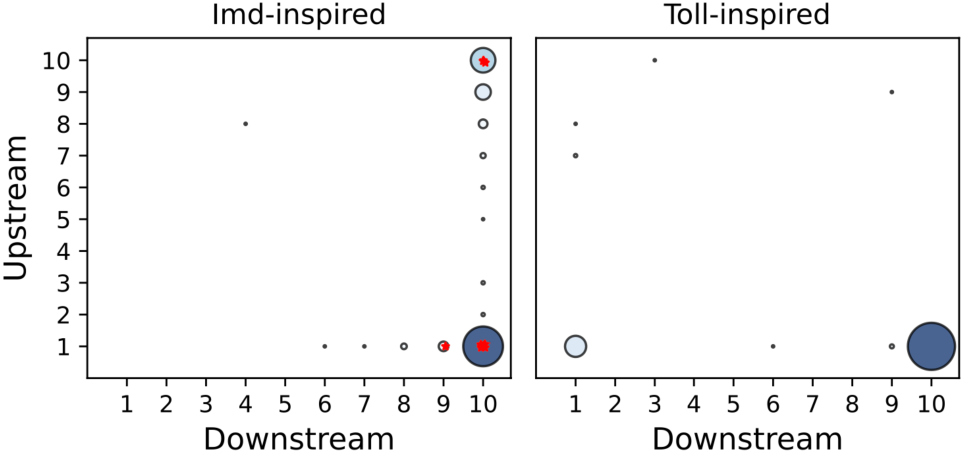
When bacterial proliferation is high, immune signaling pathways evolve either towards a downstream NFL or both NFLs. Evolution of Imd (left) and Toll signaling when bacterial proliferation rate is high (*k*_0_= 0.5). Plots show the final values of parameters controlling downstream (X-axis) and upstream (Y-axis) NFLs after 5,000 generations. Stars show simulations in which noise increases across generations. The size of the circles and the darkness of the color for every combination of NFLs are determined by the proportion of simulations that converge to that combination of NFLs.

**Fig.S7.**
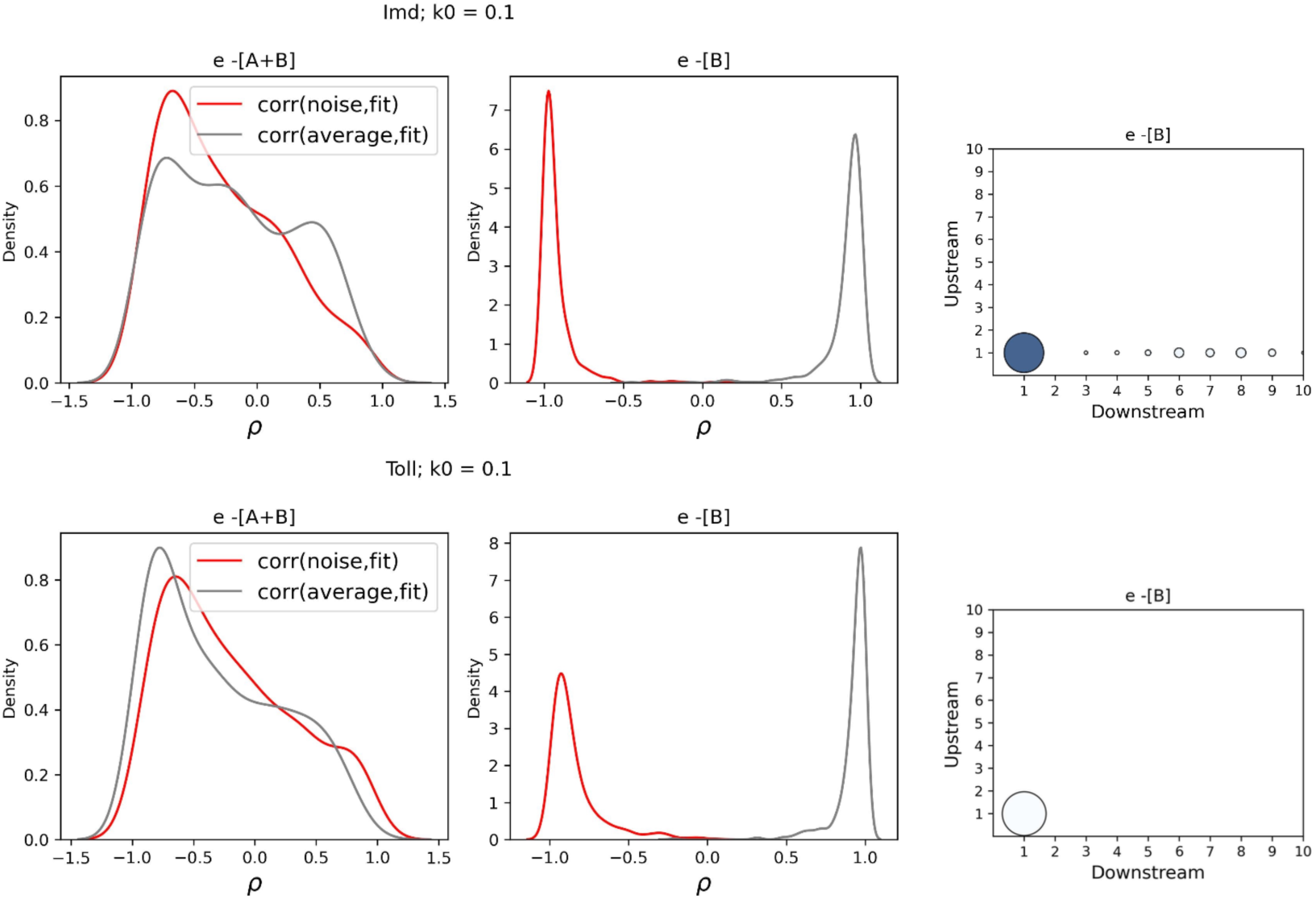
Noise only evolves when signaling pathways must reduce the cost of AMP expression. The correlation plots on the left are shown for e-[A+B] and e-[B] fitness functions. Here, we measured the Spearman correlation between noise (and average expression) and fitness across 5,000 generations of Imd and Toll evolution. Since fitness only increases during the evolution of both pathways (i.e., mutations that decrease fitness are rejected), a positive correlation with fitness indicates that the parameters (noise or average expression) is increasing across generations, whereas a negative correlation means the parameter is decreasing as the evolution continues. Here, we only see negative correlation for noise values (red plots) when we have e-[B]. On the other hand, average expression shows positive correlation when we have e-[B] to eliminate the pathogen. The results for e-[A+B] is more complicated because the signaling pathway should minimize both B and A. Minimizing A increases noise, and results in positive correlation values in some simulations when we have e-[A+B]. On the right the final combinations of upstream (Y-axis) and downstream (X-axis) NFLs are shown after 5000 generations when we have e-[B]. For all simulations presented here, we set *k*_0_ = 0.1.

**Fig.S8.**
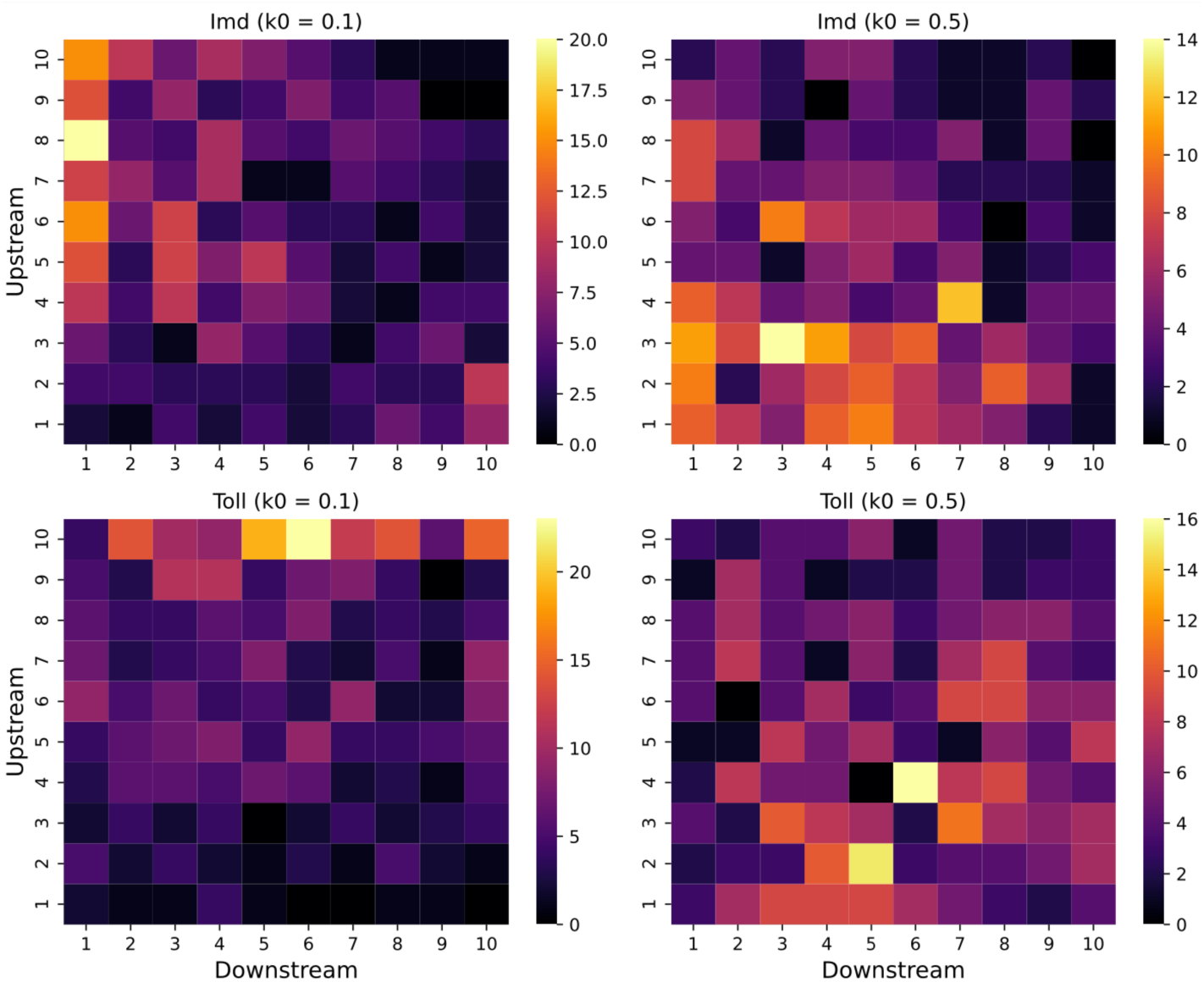
For a lower bacterial proliferation rate, both Imd and Toll signaling evolve toward an upstream-biased NFL. The evolution of a deterministic Imd (on the top) and Toll (on the bottom) signaling for two bacterial proliferation rates (*k*_0_=0.1 on the left and *k*_0_=0.5 on the right). The brighter colors show combinations of downstream and upstream NFLs that evolve more often.

