## Supplemental file for the manuscript titled Positioning of negative feedback loops within immune signaling pathways influences gene expression noise for "Positioning of negative feedback loops within immune signaling pathways influences gene expression noise"

### Gillespie simulations of the Imd and Toll models

The Imd model is a simplified (and stochastic) version of a deterministic model previously developed by [1]. Our models entail two sets of reaction rates: reaction rates pertaining to bacteria and reaction rates controlling immune signaling that responds to bacteria. If both sets are empty (i.e., all reaction rates are 0), no reactions occur. If both sets contain non-zero rates, two reactions (one for bacteria, which is either elimination or proliferation, and one for a randomly chosen protein of the immune network) occur in one step. Otherwise, one reaction occurs. The reactions are described in detail below.

After encountering a pathogen, pathogen proliferation or elimination by AMP is captured stochastically by reactions 1 and 2. The probability of bacterial proliferation $(B \rightarrow B + 1)$ is increased for a higher bacteria proliferation rate $(k_0)$ (reaction 1). The probability that bacteria are killed by AMP depends on the number of AMPs $(A_t)$ and bacteria $(B_t)$ because AMP kills the bacteria by interacting with the bacterial cell (reaction 2). The reaction rate is written in front of the reaction.

$(1)\ B \rightarrow B + 1\ (k_0 B_t)$

$(2)\ B \rightarrow B - 1\ (A_t B_t)$

Receptors are produced by NF-κB $(N_t)$, modeled by a Hill function (reaction 3). Receptors are reduced upon binding to the bacteria $(R_t B_t)$ (reaction 4) and interactions with Pirk $(P_t)$ (reaction 5) [2]. In both the Imd and Toll models, Pirk represents the upstream NFL.

$(3)\ R \rightarrow R + 1\ (\beta_1 \frac{N_t}{N_t + Z_n})$

$(4)\ R \rightarrow R - 1\ (R_t B_t)$

$(5)\ R \rightarrow R - 1\ (P_t R_t)$

The binding of bacteria forms a complex between the receptor and bacteria $(X_t)$ (reaction 6), and the complex is pulled down by Pirk (reaction 7). Formation of the complex activates NF-κB $(N_t)$ at a rate $\beta_2$ (reaction 8). NF-κB is degraded at a rate $\lambda$ (reaction 9). All proteins in the system have identical degradation rates $(\lambda)$. In reality, peptidoglycans produced by bacteria form a complex with receptors; however, for simplicity, here we assume that the bacterium itself binds to receptors.

$(6)\ X \rightarrow X + 1\ (R_t B_t)$
$(7)\ X \rightarrow X - 1\ (P_t X_t)$
$(8)\ N \rightarrow N + 1\ (\beta_2 X_t)$
$(9)\ N \rightarrow N - 1\ (\lambda N_t)$

NF-κB activates Pirk (reaction 10) and the repressosome $(S_t)$ (reaction 12) as NFLs. The repressosome complex is the downstream NFL in the Imd model.

$(10)\ P \rightarrow P + 1\ (\beta_3 \frac{N_t}{N_t + Z_n})$
$(11)\ P \rightarrow P - 1\ (\lambda P_t)$
$(12)\ S \rightarrow S + 1\ (\beta_4 \frac{N_t}{N_t + Z_n})$
$(13)\ S \rightarrow S - 1\ (\lambda S_t)$

NF-κB activates the expression of AMPs $(A_t)$. The repressosome competes with NF-κB for binding to the promoter of the AMP gene to reduce transcription. This is captured by reaction 14.

$(14)\ A \rightarrow A + 1\ (\beta_5 \frac{N_t}{N_t + Z_n + \frac{Z_n S_t}{Z_s}})$
$(15)\ A \rightarrow A - 1\ (\lambda A_t)$

The model of the Toll pathway is similar to the model of Imd. The difference is that reaction eight in the model of the Toll pathway describes the change in NF-κB outside of the nucleolus $(NO_t)$, and additional reactions describe the change in NF-κB $(N_t)$ inside the nucleolus (reactions 16 and 17):

$(16)\ N \rightarrow N + 1\ ([NO_t]\, e^{-C_t})$
$(17)\ N \rightarrow N - 1\ (\lambda N_t)$

Here $C_t$ is the concentration of Cactus, whose dynamics are described by reaction 12. Cactus prevents the flow of NF-κB to the nucleus. Therefore, if $C_t \rightarrow \infty$, $N$ does not change. Thus, Cactus acts as the downstream NFL in the Toll model [3].
Because there is no repressosome in the Toll pathway $(S_t = 0)$ and reaction 14, which describes AMP production, is reduced to:

$(18)\ A \rightarrow A\ +\ 1\ (\beta_5 \frac{N_t}{N_t + Z_n})$

$(19)\ A \rightarrow A -\ 1\ (\lambda A_t)$

The initial condition for the number of receptors is set to 10 $(R_0 = 10)$ to ensure an immune response upon infection. The initial condition for other parameters is set to zero. Below, we illustrate 10 trajectories for a fully activated Imd pathway with both NFLs present ($\beta_1, \beta_2, \beta_3, \beta_4, \beta_5 = 10,\ \lambda = 0.1, k_0 = 0.1, Z_n = Z_s = 1$).

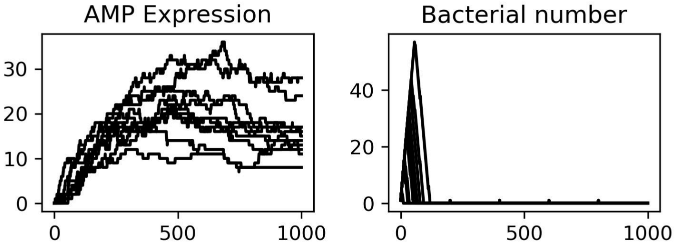


Changes in AMP expression and bacterial number

simulated using the Gillespie algorithm.

### Deterministic simulations

The deterministic solution for the system of ordinary differential equations (ODEs), formed from the reaction rates of the Imd model. Here $f(t)$ is the bacterial input.

$$\frac{dB}{dt} = f(t) + k_0 B_t - A_t B_t$$

$$\frac{dR}{dt} = \beta_1 \frac{N_t}{N_t + Z_n} - P_t R_t - R_t B_t$$

$$\frac{dX}{dt} = R_t B_t - P_t X_t$$

$$\frac{dN}{dt} = \beta_2 X_t - \lambda N_t$$

$$\frac{dP}{dt} = \beta_3 \frac{N_t}{N_t + Z_n} - \lambda P_t$$

$$\frac{dS}{dt} = \beta_4 \frac{N_t}{N_t + Z_n} - \lambda S_t$$

$$\frac{dA}{dt} = \beta_5 \frac{N_t}{N_t + Z_n + \frac{Z_n S_t}{Z_s}} - \lambda A_t$$

In the Toll model, we have an equation for N outside of the nucleus ($NO$) and another for change Inside the nucleus. Also, the change in AMP is modified according to reaction 18:

$$\frac{dNO}{dt} = \beta_2 X_t - \lambda NO_t$$

$$\frac{dN}{dt} = [NO_t]\, e^{-C_t} - \lambda N_t$$

$$\frac{dA}{dt} = \beta_5 \left(\frac{N_t}{N_t + Z_n}\right) - \lambda A_t$$
